# Homeostatic Neuronal Plasticity Alters Axon Initial Segment Actin Membrane Skeleton Periodicity

**DOI:** 10.1101/2025.10.28.685151

**Authors:** Shiju Gu, Anastasios V. Tzingounis, George Lykotrafitis

**Affiliations:** Department of Biomedical Engineering, University of Connecticut, Storrs, CT 06269, USA; Department of Physiology and Neurobiology, and University of Connecticut, Storrs, CT 06269, USA; Department of Mechanical Engineering, University of Connecticut, Storrs, CT 06269, USA

**Keywords:** Homeostatic structural neuronal plasticity, potassium channels, sodium channels, actin rings, super-resolution microscopy

## Abstract

Homeostatic plasticity allows neurons to maintain their activity around a set point, contributing to neuronal network stability over time. Increasing or decreasing neuronal activity not only regulates the levels of ion channels in neurons, but also leads to structural plasticity of the axon initial segment (AIS), the site of action potential initiation. Although previous studies have shown that activity could change the length and position of the AIS along the axon to compensate for changes in neuronal activity, it is currently unknown whether homeostatic plasticity also regulates the periodicity of the AIS actin cytoskeleton. Actin filaments, which are components of the axon plasma membrane skeleton, form periodic rings with a 190-nm spacing. Here, we examined whether the integrity of the actin ring periodicity at the AIS is plastic, depending on prolonged changes in neuronal activity. To induce AIS homeostatic plasticity, we either increased neuronal activity chronically by blocking the AIS enriched Kv7 voltage-gated potassium channels with XE991 or chronically decreased it by using the pan-Kv7 channel opener retigabine (RTG), or alternatively, the voltage-gated sodium channel blocker TTX at different neuronal maturation stages (DIV10 and DIV16). We found that prolonged exposure to XE991 relocated AIS away from soma and disrupted the periodicity of actin rings at DIV10. However, the actin ring periodicity remained unaffected at DIV16. In comparison, long-term treatment with RTG or TTX induced relocation of AIS towards the soma and reduction in AIS length, respectively. In parallel, RTG and TTX disrupted the actin ring periodicity at both developmental stages. These observations suggest that homeostatic plasticity not only changes the AIS position and length, but it could also modulate the periodicity of the AIS actin membrane skeleton, raising the possibility that the mechanical properties of the AIS are also plastic.

**Statement of significance:** Homeostatic plasticity allows neurons to maintain their activity around a set point, contributing to neuronal network stability over time. Although previous studies have shown that activity could change the length and position of the AIS along the axon to compensate for changes in neuronal activity, it is currently unknown whether homeostatic plasticity also regulates the periodicity of the AIS actin cytoskeleton. Here, using super-resolution microscopy, we show that either blocking or activating Kv7 channels could disrupt the periodicity of actin depending on the maturation stage of cultured neurons. In comparison, blocking voltage-gated sodium channels disrupted the actin ring periodicity independent of the age of neurons. These observations suggest that homeostatic plasticity also modulates the periodicity of the AIS actin cytoskeleton.

## Introduction

The axon initial segment (AIS), typically spanning 20-60 µm, is situated at the proximal axon in neurons (1). In addition to initiating action potentials (APs), the AIS serves as a selective filter that allows specific proteins and molecules to enter the axon, which ensures the polarized distribution of protein expression between the axon and other neuronal domains (2, 3). Contrary to the long-standing notion that AIS is a rigid and immobile structure, recent studies have demonstrated that the AIS can undergo changes in length and location. These adjustments enable the AIS to fine-tune neuronal excitability in response to acute or chronic stimuli, a phenomenon referred to as homeostatic plasticity. The homeostatic plasticity provides a mechanism for the AIS to dynamically modulate critical properties of APs, including threshold, amplitude, and firing frequency (4-7). For example, it has been shown that prolonged depolarization induces a distal relocation of AIS-specific proteins along the axon in various types of excitatory neurons. This relocation is associated with a decrease in excitability, as evidenced by a reduction in firing rate and an increase in threshold current required for firing APs (8, 9). In contrast, in cultured neurons from olfactory bulb, chronic depolarization (24 h) of inhibitory interneurons led to a proximal movement of the AIS towards the soma (10). These findings suggest that, during plasticity, the movement of the AIS along the axon varies among different types of neurons.

KCNQ (Kv7) channels are integral in regulating neuronal excitability, primarily through their suppression of burst and repetitive AP firing as the molecular correlates of the M-current, a voltage-activated non-inactivating potassium channels resident in AIS (12, 13). The distinct functional significance of Kv7 channels in inhibiting excitability highlights their critical role in preventing seizures and epilepsy (14, 15). Several studies have have also implicated Kv7 channels in neuronal plasticity. For example, chronic blockade (48 h) of neuronal activity led to an increase in the AP firing rate and a decrease in Kv7 current (16). In another study, it was shown that acute treatment (<1 hr) with XE991 increased neuronal excitability. However, with sustained application of XE991, a gradual reduction in intrinsic excitability was observed. Furthermore, after prolonged exposure (48 hrs) to XE991, several AIS components, including ankyrin G (ankG), Nav channels, and Kv7 channels, were observed to relocate significantly towards a more distal position along the axon, without changing their axonal lengths (17). In contrast to XE991, retigabine (RTG) holds Kv7 channels open and hyperpolarizes the plasma membrane, thereby limiting generations of Aps (18) reducing neuronal excitability. However, following chronic exposure of retigabine (4-48 hrs), the threshold current and spontaneous firing rate normalized and this was accompanied by a proximal relocation of ankG and Kv7 channels towards the soma (19).

Neuronal plasticity is also induced by blockage of Nav channels. During the upstrong of an AP, Nav channels at AIS open, allowing influx of sodium ions and membrane depolarization. Nav channels are anchored to the AIS through their interaction with the scaffolding protein AnkG (20, 21), and this interaction is regulated by the protein kinase CK2 (22). The density and biophysical properties of Nav channels are key factors of the AP threshold. For example, in cortical pyramidal neurons, the density of Nav channels at the AIS has been reported to be approximately 50 times higher than that in the proximal dendrite (20). Previous work found that, following long-term incubation with tetrodotoxin (TTX), a potent Nav channel blocker, the AP firing rate of hippocampal neurons increased (23-25). Additionally prolonged incubation (48 hrs) with TTX caused HCN1 channels to translocate proximally towards the soma in CA1 pyramidal neurons (26). In another study, after a 24 hr treatment with TTX in rat olfactory bulb dopaminergic neurons, the length of the AnkG signal and consequently of the AIS decreased, whereas the starting location along the axon remained unchanged (10). A more recent study reported that prolonged application (48 hrs) of TTX did not affect AnkG length, but it reduced ankG immunodensity (27). The observed differences between these findings may be attributed to differences in cell types studied.

Actin filaments are a major component of the cytoskeleton providing cells with essential mechanical support, helping to maintain cell morphology. Actin filaments are involved in various cellular processes, including migration and division (28). Super-resolution microscopy has revealed that the axon membrane skeleton comprises actin rings periodically spaced along the axon at a distance of approximately 190 nm and connected via longitudinal spectrin tetramers (29, 30). AnkG functions as a molecular bridge, facilitating the connection between the actin-spectrin network and membrane proteins to the internal microtubules (31). The axonal periodic actin assembly initiates at days in vitro (DIV) 2 and becomes apparent by DIV6, which coincides with the day that the actin capping protein adducin adopts a periodic pattern (29, 30, 32). Results from a prior study suggest that these periodic actin rings act as barriers, limiting diffusion of small-conductance calcium-activated potassium (SK) channels at DIV6 (33). Furthermore, at DIV14 and DIV18, SK channels remain immobilized at the AIS following a 0.5-h treatment with Swinholide A (SwinA), a potent actin-disrupting drug (33, 34). This finding indicates that high concentration of components within the AIS might generate a crowding effect that possibly restricts diffusion of SK channels independently of the presence of actin ring barriers (35, 36). Despite extensive investigation into homeostatic structural plasticity, the integrity of the actin rings at AIS during short-(1-4 hrs) and long-term (>24 hrs) neuronal plasticity is unclear.

Here, we induced homeostatic neuronal plasticity by treating cultured hippocampal neurons with the pan-Kv7 channel blocker XE991, the Kv7 channel opener RTG, and the Nav channel blocker TTX at DIV10 and DIV16, for various time periods. In addition to systematically measuring both the position and length of the AIS, we investigated the periodicity of actin rings at the AIS under each of these treatment conditions. Our findings indicate that prolonged incubation with XE991 progressively induced a distal relocation of the AIS. This treatment was associated with a gradual decrease of the degree of periodicity of actin rings in the AIS at DIV10, while no significant change was detected at DIV16. In contrast to the effects of XE991, blocking Nav channels with TTX led to a shortening of the AIS length, without changing its starting position. Meanwhile, prolonged exposure to RTG caused a proximal relocation of the AIS towards the soma. Notably, after prolonged treatment with TTX and RTG (48 hrs), the degree of actin ring periodicity significantly decreased in both groups at both DIV10 and DIV16. In conclusion, our study demonstrates the plasticity of actin rings at the AIS following either increased or decreased prolonged neuronal activity

## Materials and methods

### Neuron Culture

Hippocampal tissues from embryonic day 18 rats were purchased from BrainBits (SDEHPS; Springfield, IL, USA). The tissues were dissociated into single neurons using a 0.05% Trypsin-EDTA solution (15400054; Thermo Fisher Scientific, Waltham, MA, USA). Subsequently, the dissociated neurons were plated onto coverslips (12-545-83; Fisher Scientific International, Inc., Hampton, NH, USA), which had been pre-coated with poly-D-lysine (P0899; MilliporeSigma, Burlington, MA, USA), at a final density of 50,000 neurons per coverslip. The culture medium for these neurons consisted of Neurobasal Medium (21103049; Thermo Fisher Scientific), B-27 Supplement (17504044; Thermo Fisher Scientific), Penicillin-Streptomycin (15140122; Thermo Fisher Scientific), and GlutaMax Supplement (35050061; Thermo Fisher Scientific). We changed half of the culture medium every three days. The neurons were incubated at 37 °C with 5% CO_2_ until they were used.

### Immunocytochemistry

Neurons were treated with 10 µM XE991, 10 µM RTG, 1 µM TTX or 0.1% (v/v) Dimethylsulfoxide (DMSO) vehicle for a designated time period prior to staining. Following this treatment, the cells were quickly washed three times with DPBS (14190144; Thermo Fisher Scientific). The neurons were then fixed with 4% paraformaldehyde (15710; Electron Microscopy Sciences; Hatfield, PA, USA) for 15 min at room temperature (RT). To quench the background fluorescence caused by paraformaldehyde fixation, the neurons were washed three times and incubated with 50 Mm NH_4_Cl (A9434; MilliporeSigma) for an additional 10 min. The neurons were subsequently washed three times and treated with 0.1% Triton X-100 (93443; MilliporeSigma) for 5 min to permeabilize the cell membrane. After three washes, the neurons were labelled with mouse ankyrin G antibody (1:200; sc-12719; Santa Cruz Biotechnology, Dallas, TX, USA) in pre-block buffer (5% normal goat serum in PBS) and incubated overnight at 4 °C. The following day, the neurons were washed three times and incubated with goat anti-mouse abberior STAR ORANGE secondary antibody (1:200; STORANGE-1001-500UG; Abberior; Göttingen, Germany) in pre-block buffer to label the ankyrin G antibody. Meanwhile, to label actin filaments, we also added abberior STAR RED phalloidin conjugation (30Nm; STRED-0100-20UG; Abberior) for 2 hrs at RT in the dark. Finally, the neurons were washed six times and mounted on glass slides using Prolong Diamond Antifade Mountant (P36965; Thermo Fisher Scientific). The slides were allowed to dry at RT in the dark for 24 hrs and then imaged immediately.

### Image Acquisition and Analysis

All AnkG images were acquired using a laser-scanning Leica SP8 confocal microscope (Leica Microsystem, Wetzlar, Germany). A 561-nm laser was employed to excite the fluorescence of the abberior STAR ORANGE molecules conjugated to AnkG. Z-stack images were subsequently collected and we performed maximum intensity projection for the stack images. The confocal image acquisition procedure was conducted utilizing the LAS X software (Leica Microsystem).

For the super-resolution imaging of actin rings at the AIS, a stimulated emission depletion (STED) microscope was utilized. Fixed neurons samples were imaged using a 100X oil immersion lens. A 595-nm excitation pulsed laser was used to excite the fluorescence from abberior STAR ORANGE molecules, thereby identifying the location of the AnkG proteins. In the same region, both a 640-nm excitation pulsed laser and a 775-nm depletion laser were applied to scan the STAR RED molecules conjugated to the actin filaments. Image acquisition with STED was performed using the Abberior Imspector (Abberior) software. The pixel size for all scanned actin images was consistently set at 30 nm.

All images were analyzed using ImageJ software (37). To evaluate the location and length of AnkG, we drew a line along AnkG signal, extending from the soma to the point where the signal disappeared. We then plotted the signal intensity and transferred the raw signal values to Excel spreadsheet. Each signal intensity was normalized to a range of 0 to 1 by dividing by the maximum value. Based on this normalized fluorescence intensity plot, we determined the start and end position of AnkG. For the proximal axon, the start of the AnkG signal was identified when the normalized value exceeded 0.33 and persisted for a minimum of 5 µm. For the distal axon, the end of the AnkG signal was identified when the normalized value fell below 0.33 and persisted for a minimum of 2 µm. The length of AnkG was defined as the distance between these identified start and end positions. Notably, the start position and length of AnkG determined through this method aligned well with observations made visually (8, 38).

To quantify the degree of periodicity of actin rings at the AIS, we first drew a 3 µm-long line at a random location along AIS on an actin STED image. We then extracted the fluorescence intensity values along this line and transferred them to an Excel spreadsheet. The autocorrelation was computed using the following equation:

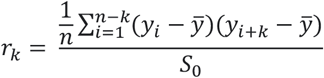

Here, *k* is the lag, 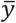 denotes the mean of a time series: *y*_1_, …,*y*_*n*_, *S*_0_ represents the variance of the time series. *r*_*k*_ is the autocorrelation value at lag *k*.

It has been found that the distance between adjacent axonal actin rings is approximately 190 nm (29). Hence, to calculate the autocorrelation amplitude, we subtracted the average value of the first two valleys (at 95 nm and 285 nm) from the value of the first peak (at 190 nm). If the autocorrelation curve exhibited more than two 190 nm periodic peaks, we categorized the actin rings at this region as periodic; otherwise, they were classified as non-periodic (39).

### Statistical Analysis

Statistical analysis were conducted using GraphPad Prism (GraphPad Software, Inc., San Diego, CA, USA). To compute the results, we either performed non-parametric one-way ANOVA test or non-parametric Mann-Whitney test, as appropriate. A result was consider statistically significant if the p-value was less than 0.05. The significance levels are denoted as follows: * *P* < 0.05, ** *P* < 0.01, *** *P* < 0.001, \*\*\*\**P* < 0.0001.

## Results

The plasma membrane skeleton of the AIS is formed by periodic actin rings spaced every 190 nm and interconnected by spectrin tetramers (29). In this work, we aim to determine the structural integrity of axon periodic membrane skeleton (APMS) in the AIS following induction of homeostatic structural plasticity. To identify the location and length of the AIS, we tracked AnkG using confocal microscopy. As the organizing protein in the AIS, AnkG clusters several membrane proteins, including cell adhesion molecules such as neurofascin 186 (NF-186), NrCAM, and the Nav and Kv channels (40). Specifically, to evaluate location and length of AIS, we used hippocampal neurons and labelled them with AnkG antibody and the abberior STAR ORANGE fluorophore. After acquiring images of AnkG, we drew a line along the AnkG fluorescent signal (red line in Figure S1A) and normalized the signal intensity (red plot in Figure S1B) to determine the start and end positions of AnkG (magenta dash line in Figure S1B). Length of AnkG fluorescent signal was then calculated, as shown in Figure S1B. Further details are provided in the Methods section. Next, we examined neuronal plasticity in both immature and mature neurons. We designated DIV10 for immature neurons and DIV16 for mature ones. We selected DIV10 because at this time point the periodic actin rings at the AIS have already become prominent (29, 30, 32), while several AIS molecules, such as neurofascin, Nav channels, and end-binding (EB) protein3, have not yet reached peak levels at the AIS (35, 41, 42). This leads to a less crowded AIS at DIV10 in comparison to DIV16, a time point widely accepted as the stage of full neuronal maturation,(17, 43-45) (33, 46, 47) Add here the recent Science paper

We first examined if DMSO, the solvent used to dissolve all compounds, could influence the distribution of AnkG along the axon. Neurons at DIV8 and DIV14 were treated with DMSO at a final concentration of 0.1% (v/v) for 48 hrs and were then fixed at DIV10 and DIV16, respectively. At both DIVs, compared with the data from the non-DMSO treated group, we found that DMSO did not induce any significant changes in either the starting position of AnkG (Figure S1C; DIV10 No drug: *Start* = 6.91 ± 3.63 *μm*; DIV10 DMSO: *Start* = 7.13 ± 3.83 *μm, P* = 0.9957 ; DIV16 No drug: *Start* = 5.16 ± 2.31 *μm* ; DIV16 DMSO: *Start* = 5.12 ± 3.31 *μm, P* = 0.5718), or the length of AnkG signals along the axon (Figure S1D; DIV10 No drug: *Length* = 29.58 ± 6.85 *μm* ; DIV10 DMSO: *Length* = 29.33 ± 10.32 *μm, P* = 0.7729; DIV16 No drug: *Length* = 30.36 ± 7.77 *μm*; DIV16 DMSO: *Length* = 30.94 ± 8.60 *μm, P* = 0.8245). These results suggest that DMSO alone does not affect the distribution of AnkG at the AIS. Therefore, we used neurons treated with DMSO as our control group in the following experiments.

To explore how changes in neuronal excitability influence location of the AIS with respect to the soma and its length, we treated neurons with the pan-Kv7 channel blockerXE991, the pan-Kv7 channel opener RTG, or the Nav channel blocker TTX. In particular, we treated cultured neurons with 10 µM XE991, 10 µM RTG, or 1 µM TTX for 12, 24, 36, or 48 hrs prior to fixation. Neurons treated with DMSO for 48 hrs were used as control. Representative AnkG fluorescent signals and their corresponding normalized fluorescence intensity plot from the control group are shown in Figure 1A(i) and Figure 1B(i), respectively. At DIV10, we found that XE991 application induced a distal shift in the entire AnkG signal along the axon, as illustrated in Figure 1A(ii) and Figure 1B(ii). After measuring the starting positions of AnkG signal in both the control and XE991 treated groups, we applied non-parametric one-way ANOVA test and a significant increase was found after 24-hrs treatment with XE991 [Figure 1E(i); DMSO: *Start* = 5.88 ± 1.94 *μm*; XE991_24 h: *Start* = 8.52 ± 3.48 *μm, P* = 0.0452], while the length of the AnkG signal along the axon remained unchanged [Figure 1E(ii)]. Our result suggests that prolonged blockade of Kv7 channels with XE991, which increases neuronal excitability, prompts the neuron to initiate a homeostatic plasticity, which gradually relocates its AIS away from the soma, thereby reducing excitability. These findings are in agreement with previous work (17).

**Figure 1.**
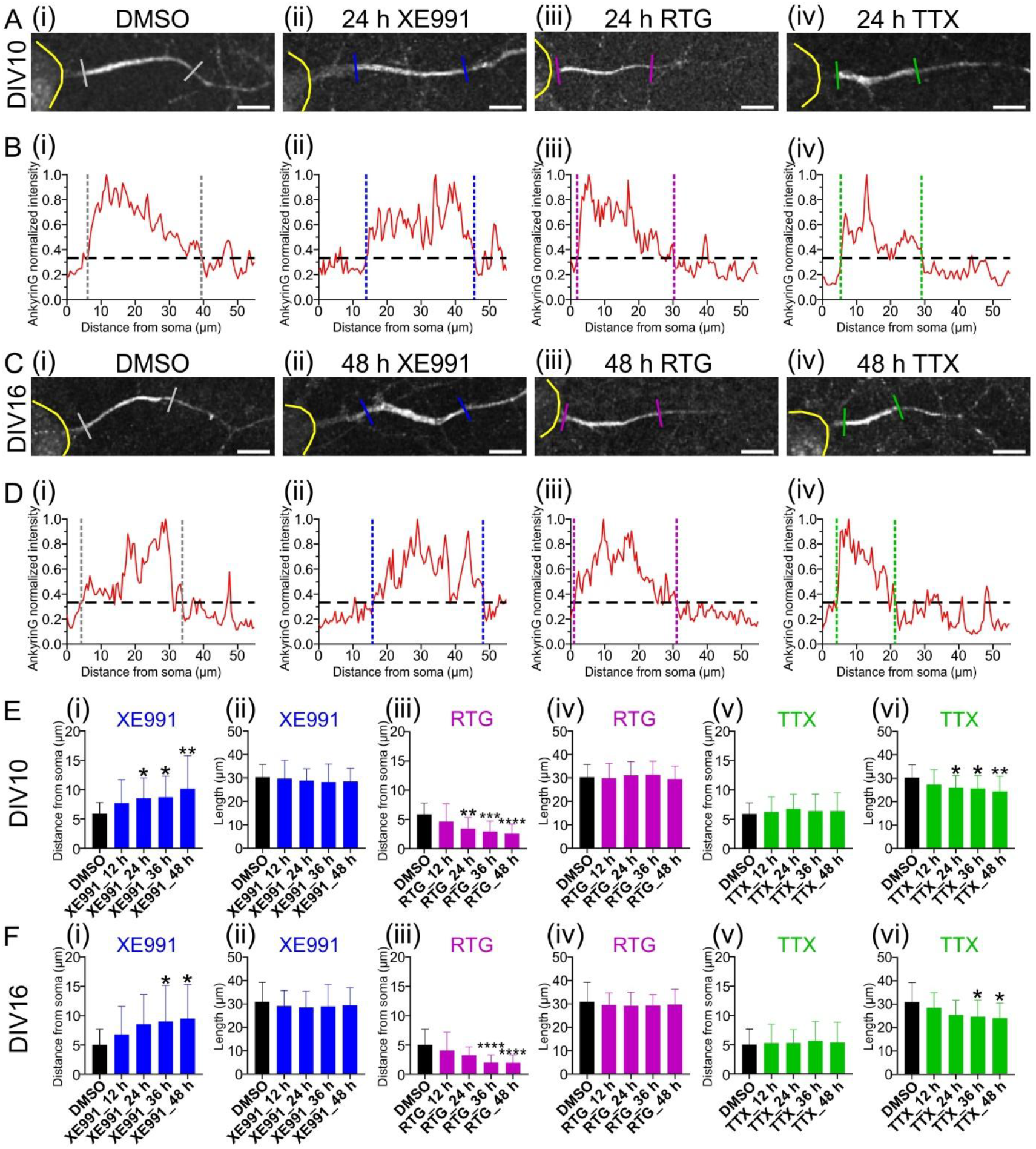
Effects of XE991, RTG, and TTX treatments on ankG distribution in cultured hippocampal neurons at DIV10 and DIV16. *(A)* Representative confocal microscopy images of DIV10 neurons stained for AnkG after 48-h treatment with DMSO (control), or 24-hrs treatment with XE991, RTG, or TTX. Scale bar, 10 µm. *(B)* Plots showing the normalized fluorescence intensity of ankG along the axon, corresponding to the images shown in *(A). (C)* Representative confocal images of DIV16 neurons stained for AnkG after 48-h treatment with DMSO, XE991, RTG, or TTX. Scale bar, 10 µm. *(D)* Plots presenting the normalized fluorescence intensity of ankG along the axon, corresponding to the images shown in *(C). (E)* Plots showing the starting position and length of ankG along the axon at DIV10 after treatment with XE991 *(i-ii)*, RTG *(iii-iv)*, or TTX *(v-vi)*. Sample size: *(i)* and *(ii)*: DMSO: n = 21; XE991_12 h: n = 20; XE991_24 h: n = 20; XE991_36 h: n = 20; XE991_48 h: n = 20. *(iii)* and *(iv)*: DMSO: n = 21; RTG_12 h: n = 25; RTG_24 h: n = 25; RTG_36 h: n = 25; RTG_48 h: n = 25. *(v)* and *(vi)*: DMSO: n = 21; TTX_12 h: n = 24; TTX_24 h: n = 22; TTX_36 h: n = 22; TTX_48 h: n = 24. *(F)* Plots showing the start position and length of ankG at DIV16 following the same treatments as applied at DIV10 in *(E)*. Sample size: *(i)* and *(ii)*: DMSO: n = 21; XE991_12 h: n = 20; XE991_24 h: n = 20; XE991_36 h: n = 20; XE991_48 h: n = 20. *(iii)* and *(iv)*: DMSO: n = 21; RTG_12 h: n = 25; RTG_24 h: n = 25; RTG_36 h: n = 25; RTG_48 h: n = 25. (v) and *(vi)*: DMSO: n = 21; TTX_12 h: n = 23; TTX_24 h: n = 23; TTX_36 h: n = 23; TTX_48 h: n = 23. \**P* < 0.05, \*\**P* < 0.01, \*\*\**P* < 0.001, \*\*\*\**P* < 0.0001 (non-parametric one-way ANOVA test). All plots represent the mean ± standard deviation (SD).

As a second test, we used RTG, which in contrast to XE991, led to a shift of the ankG signal towards the soma after 24-h of incubation [Figure 1E(iii); DMSO: *Start* = 5.88 ± 1.94 *μm*; RTG_24 h: *Start* = 3.45 ± 1.86 *μm, P* = 0.0038], without affecting its length along the axon [Figure 1E(iv)]. This result, which is in agreement with prior experimental results(19), suggests that as RTG keeps the Kv7 channels open, the membrane potential decreases, leading to reduced excitability. To compensate and restore excitability towards normal levels, neurons gradually shift the AIS closer to the soma. Finally, we repeated our tests using TTX. We found that, unlike XE991 and RTG, TTX did not affect the start position of AnkG [Figure 1E(v)], but significantly reduced the overall length of the ankG signal after a 24-h treatment [Figure 1E(vi); DMSO: *Length* = 30.32 ± 5.40 *μm*; TTX _24 h: *Length* = 25.95 ± 5.12 *μm, P* = 0.0417]. It’s noteworthy that our results regarding TTX induced plasticity align with findings from a prior study (10).

To determine if plasticity in mature neurons produces different results compared to immature neurons, we replicated our experiments at DIV16. We noticed that the results at DIV16 are similar to the ones found at DIV10. Following a 36-h treatment with XE991, we observed a significant shift of the AIS away from the soma [Figure 1F(i); DMSO: *Start* = 5.02 ± 2.65 *μm* ; XE991_36 h: *Start* = 9.03 ± 6.12 *μm, P* = 0.0463]. Conversely, after 36-h of RTG exposure, AnkG and consequently the AIS noticeably relocated closer to the soma [Figure 1F(iii); DMSO: *Start* = 5.02 ± 2.65 *μm*; RTG_36 h: *Start* = 2.00 ± 1.32 *μm, P* < 0.0001]. Neither drug influenced the length of the AnkG signal along the axon [Figure 1F(ii) and Figure 1F(iv)]. In contrast, a 36-h treatment with TTX reduced the length of the AIS [Figure 1F(vi); DMSO: *Length* = 30.96 ± 8.28 *μm*; TTX _36 h: *Length* = 24.74 ± 7.04 *μm, P* = 0.0364], without altering its starting position [Figure 1F(v)]. These results suggest that long-term homeostatic structural plasticity affects AIS location and length along the axon similarly in both mature (DIV16) and immature (DIV10) neurons. Furthermore, we also examined short-term homeostatic structural plasticity by treating neurons with XE991, RTG, or TTX for 1 and 4 h at both DIV10 and DIV16. These time points were selected based on previous studies, which have shown that homeostatic compensatory mechanisms can be triggered then (17, 19). After treating neurons with the XE991, RTG, or TTX for 1 and 4 h, we observed no significant alterations of the AIS under these conditions (Figure S2).

Up to this point, we have reaffirmed prior observations regarding the influence of XE991 and RTG on AnkG positioning and AIS length. Additionally, we re-examined the impact of TTX on AnkG distribution along the axon. Next, we aimed to explore whether the integrity of periodic structure of the actin rings is related to AIS movement when exposed to the three drugs used to stimulate neuronal homeostatic plasticity at DIV10 and DIV16. To achieve this, we utilized STED super-resolution microscopy. In particular, we used phalloidin STAR RED conjugation to specifically label actin filaments, as phalloidin could bind to the interface between F-actin subunits (48). We first imaged the AnkG signal. A 595-nm excitation pulsed laser was used to excite the fluorescence from abberior STAR ORANGE molecules, thereby identifying the location of the AnkG proteins. To locate the AIS, we employed an excitation laser to excite the fluorescence from abberior STAR ORANGE, conjugated to AnkG, as detailed in the Methods section. Subsequently, we scanned the actin filaments in the same region using STED (Figure 2A). To quantify the periodicity of actin rings, we selected a random region of the AIS, drew a line across this region, and performed autocorrelation analysis, as shown in Figure 2B. We calculated the autocorrelation amplitude for each selected region, as well as the fraction of regions that exhibited a periodic signal, in accordance with the procedures detailed in the Methods section.

**Figure 2.**
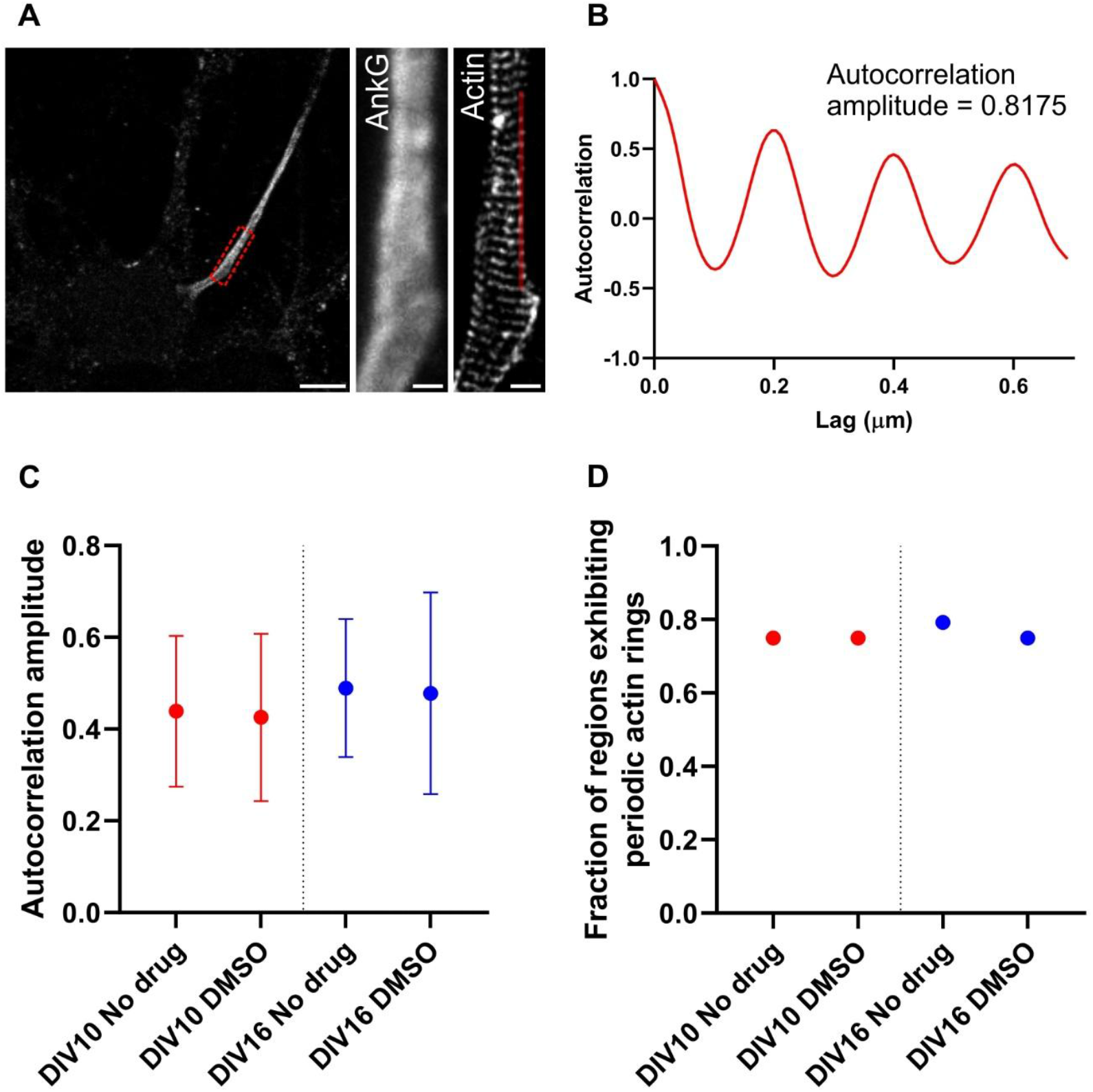
DMSO Treatment does not alter the periodicity of actin rings at the AIS. *(A) Left*: Immunolabeling of AnkG in cultured neurons. *Middle*: Enlarged image of AnkG, corresponding to the red boxed region in the *left. Right*: STED image of actin rings in the same region. Scale bar, *left*: 10 µm, *middle*: 0.5 µm, *right*: 0.5 µm. *(B)* Autocorrelation plot derived from the red line region indicated in the *(A) Right. (C)* Comparison of the autocorrelation amplitude of actin rings between non-drug treatment and DMSO treatment groups at DIV 10 and 16. Sample size: n = 24 for each treatment group. Non-parametric Mann-Whitney tests were performed. The plots represent the mean ± SD. *(D)* Fraction of regions exhibiting periodic actin rings in non-drug treatment and DMSO treatment groups at DIV10 and DIV16.

We first evaluated the effects of DMSO on actin rings periodicity because DMSO was used as solvent for the plasticity inducing drugs. Our results at both time points, DIV10 and DIV16, indicated that a 48-hr application of DMSO did not significantly alter the autocorrelation amplitude at the AIS (Figure 2C; DIV10 No drug: *Amplitude* = 0.44 ± 0.16; DIV10 DMSO: *Amplitude* = 0.43 ± 0.18, *P* = 0.9268; DIV16 No drug: *Amplitude* = 0.49 ± 0.15 ; DIV16 DMSO: *Amplitude* = 0.48 ± 0.22, *P* = 0.9106). Moreover, we observed that the fraction of regions exhibiting periodic actin rings remained consistently high, regardless of DMSO treatment (Figure 2D; DIV10 No drug: *Fraction* = 0.75; DIV10 DMSO: *Fraction* = 0.75; DIV16 No drug: *Fraction* = 0.79; DIV16 DMSO: *Fraction* = 0.75). These findings suggested that DMSO treatment does not disrupt periodicity and integrity of the actin rings at the AIS. Hence, we used neurons treated with DMSO as control in our subsequent tests.

We found that at DIV10, neurons in the control group have developed periodic actin rings along the AIS, as shown in Figure 3A(i), with the corresponding autocorrelation plot presented in Figure 3B(i). However, prolonged incubation with XE991, RTG, and TTX markedly reduced the degree of periodicity of actin rings at the AIS [Figure 3A(ii)-(iv) and Figure 3B(ii)-(iv)], as illustrated by a significant decline in autocorrelation amplitude. Specifically, a 24-hr incubation with XE991 resulted in a notable reduction in the autocorrelation amplitude [Figure 3E(i); DMSO: *Amplitude* = 0.42 ± 0.18 ; XE991_24 h: *Amplitude* = 0.28 ± 0.19, *P* = 0.0273]. Similarly, after a 12-hr treatment with RTG or TTX, we recorded a significant decrease in the autocorrelation amplitude [Figure 3E(iii); DMSO: *Amplitude* = 0.42 ± 0.18 ; RTG_12 h: *Amplitude* = 0.24 ± 0.14, *P* = 0.0126 and Figure 3E(v); DMSO: *Amplitude* = 0.42 ± 0.18; TTX_12 h: *Amplitude* = 0.28 ± 0.19, *P* = 0.0455]. In addition, the fraction of regions exhibiting periodic structures also diminished [Figure 3E(ii),(iv),(vi)]. These findings suggest that at DIV10 during the homeostatic plasticity triggered by treating neurons with XE991, RTG, or TTX, there’s a partial disruption in the actin rings, possibly facilitating relocation of AIS proteins and promoting endocytosis/exocytosis (Letter… reference..).

**Figure 3.**
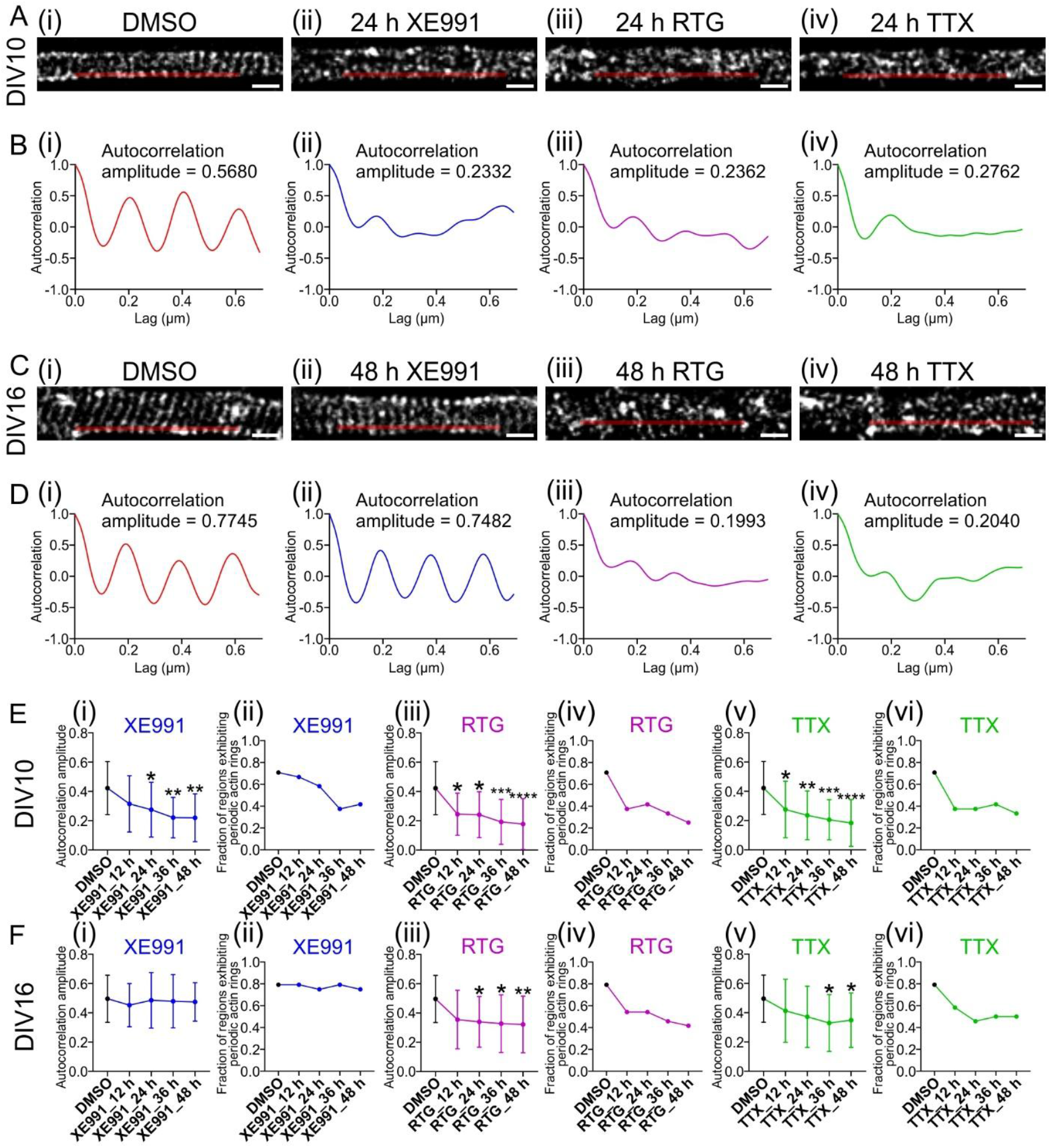
Long-term incubation of XE991, RTG or TTX differentially affects the structure of actin rings at the AIS. *(A)* Representative STED microscopy images of actin rings at the AIS of DIV10 neurons after 48-h treatment of DMSO (control), or 24-h treatment with XE991, RTG, or TTX. Scale bar, 0.5 µm. *(B)* Autocorrelation plots derived from the red line regions in *(A). (C)* Representative STED images of actin rings at the AIS of DIV16 neurons after 48-h treatment with DMSO, XE991, RTG, or TTX. Scale bar, 0.5 µm. *(D)* Autocorrelation plots derived from the red line regions in *(C). (E)* Plots showing autocorrelation amplitude and fraction of regions exhibiting periodic actin rings at DIV10 following treatment with XE991 *(i-ii)*, RTG *(iii-iv)* or TTX *(v-vi)*. Sample size: n = 24 for each treatment group. *(F)* Plots showing autocorrelation amplitude and fraction of regions exhibiting periodic actin rings at DIV16 following the same treatments as applied at DIV10 in *(E)*. Sample size: n = 24 for each treatment group. \**P* < 0.05, \*\**P* < 0.01, \*\*\**P* < 0.001, \*\*\*\**P* < 0.0001 (non-parametric one-way ANOVA test). All autocorrelation amplitude plots represent the mean ± SD.

Considering that the AIS forms a more robust assembly and the potential membrane crowding in mature neurons as mentioned previously (35, 36), we repeated our experiments at DIV16. Interestingly, treatment with XE991 did not impact the actin ring structures, as indicated by stable autocorrelation amplitude and a consistently high fraction of regions displaying periodic rings [Figure 3C(i)-(ii), Figure 3D(i)-(ii), and Figure 3F(i)-(ii)]. We hypothesize that at DIV16, when neurons are mature, the plasma membrane crowding at AIS may either hinder the actin disintegration process or inhibit the movement of fragmented actin filaments during homeostatic structural plasticity. This result is agreement with a previous study, where application of the actin-disrupting drug SwinA, was shown to restore SK channel diffusion at DIV10 at AIS. However, this restoration was not observed at DIV14 and DIV18, a stage when neurons achieved full maturation and the membrane at AIS may became crowded (33).

In contrast, neurons treated with RTG and TTX at DIV16 displayed diminished degree of periodicity of actin rings [Figure 3C(iii)-(iv) and Figure 3D(iii)-(iv)]. Specifically, following a 24-hr incubation with RTG, we recorded a significant decrease in autocorrelation amplitude [Figure 3F(iii); DMSO: *Amplitude* = 0.50 ± 0.16; RTG_24 h: *Amplitude* = 0.34 ± 0.17, *P* = 0.0128], and a similar finding was observed after a 36-hrs of TTX treatment [Figure 3F(v); DMSO: *Amplitude* = 0.50 ± 0.16 ; TTX_36 h: *Amplitude* = 0.33 ± 0.19, *P* = 0.0114]. Following prolonged incubation, both drugs greatly reduced the fraction of regions exhibiting periodic actin rings in the AIS [Figure 3F(iv) and 3F(vi)]. The reduced autocorrelation amplitude and periodic fraction, implying compromised actin rings periodicity, suggests that treatment with TTX and RTG may either diminish the potential crowded environment at the AIS or trigger responses that enable actin dismantlement and movement.

Additionally, in order to test the impact of short-term plasticity process on the periodicity of actin rings, we conducted further experiments applying 1- and 4-hrs treatments with XE991, RTG, or TTX at both DIV10 and DIV16, and found no significant alterations to actin rings at the AIS in these conditions (Figure S3). This results suggest that short-term treatment with these drugs is insufficient to significantly effect the integrity of the periodic actin rings at AIS.

## Discussion

While relocation of AIS proteins and changes in neuronal excitability during homeostatic structural neuronal plasticity have been explored, structural changes of the membrane skeleton in such processes remain largely unexplored. In this study, we investigated how stable is the periodicity of actin rings along the AIS when homeostatic plasticity was induced by XE991, RTG, or TTX; compounds that target channels enriched at the AIS. Our results showed that sustained membrane depolarization, achieved by blocking Kv7 channels with XE991, prompted a distal shift of AIS along the axon. Concurrently, at DIV10, after exposing to XE991 for 24 hrs, the periodicity of actin rings at the AIS notably declined. However, actin periodicity remained at a high level after long-term exposure to XE991 at DIV16 up to 48 hrs. In contrast, chronic hyperpolarization induced by either the Kv7 channel opener RTG or the Nav channel blocker TTX resulted in the movement of AIS closer to the soma and a reduction in the length of the AIS, respectively. At DIV10, 12-hr incubation with either RTG or TTX significantly reduced the periodicity of actin rings. Similarly, at DIV16, this effect was observed after 24-hr incubation with RTG and 36-h incubation with TTX.

Relocation of the AIS away from soma or a reduction in its length is indicative of a decrease in neuronal excitability (8, 49). In our work, blocking Kv7 channels with XE991, which causes membrane depolarization, would lead to an increase in neuronal excitability. In response to this heightened excitability, the AIS strategically repositioned itself further from the soma, serving as a negative-feedback compensatory mechanism. Conversely, when Kv7 channels were held open by RTG, leading to a reduction in excitability, the AIS shifted closer to the soma to counterbalance the diminished excitability. However, in our study, after blocking Nav channels with TTX, which decreases excitability, rather than extending AIS length to compensate for the reduction in neuronal excitability, the length of the ankG signal contracted. What could be the underlying mechanism for this response? As the density of Nav channels present at the AIS is very high (50) and this density likely remains unchanged during plasticity (8, 49), blocking these densely packed Nav channels with TTX would dampening neuronal excitability, not necessarily substantially affecting the resting membrane potential. Due to the extensive blockade of Nav channels at the AIS by TTX, we speculate that it might prevent recruitment of voltage-gated calcium channels (ref Science paper), which may explain the observed reduction in AnkG signal length. Nevertheless, the same treatment with TTX had the opposite effect on AIS length in excitatory cortical layer 2/3 neurons, implying that the influence of neuronal plasticity on AIS is cell-type dependent (51). The precise mechanisms behind these contrasting results require further investigation.

In a previous study, it was found that the periodicity of actin rings in the AIS persisted even after a 3-hr treatment with SwinA, which causes actin filament degradation, in mature neurons (43). The question then arises: how do actin rings remain intact despite being exposed to potent actin-disrupting drugs for such duration? We hypothesize that this may be attributed to the dense molecular environment at the AIS. By DIV16, several AIS-specific proteins or molecules, including AnkG, β4-spectrin, Neurofascin, NrCAM, EB3, and others, have been shown to reach maximal accumulation and assemble into the actin-based periodic cytoskeleton (32, 41, 42). Together, these molecules create a highly crowded environment at the AIS. As a result, it is possible that actin rings remain in place and maintain a periodic pattern, even their assembly may be partially disrupted.

Here, we observed that treating neurons with XE991 up to 48 hrs at DIV16 did not diminish the integrity of actin rings at the AIS. This finding aligns with the previous hypothesis that there might be a crowding effect at AIS in mature neurons. Despite the possible undergoing fragmentation, the actin filaments retained their position and continued to display a periodic arrangement. However, why does prolonged exposure to TTX and RTG result in the observed breakdown of actin rings at DIV16. It has been previously reported that Nav channels are essential for the formation of nascent AIS in spinal motor neurons (52) and that remodeling of actin-spectin filaments depends on calcium levels in cultured excitatory neurons (Science paper). Therefore, either silencing neurons or preventing their spontaneous firing activity also limits calcium influx, thereby reducing calcium dependent signaling pathways. These findings further highlight the critical role of Nav channels in AIS formation and stability. In our study, we speculate that after long-term blocking Nav channels with TTX at DIV16, clustering of multiple AIS membrane proteins was reduced, leading to a less densely populated AIS. Consequently, the disassembled actin segments were able to move away from their original positions resulting in non-periodic actin rings under microscopic observation. However, the precise mechanism requires further investigation.

Similarly, XE991 has been reported to facilitate the influx of calcium ions. Moreover, the calcium influx and membrane depolarization induced by XE991 application is counteracted by TTX (57). Based on these findings, we propose that in our our work, exposure to XE991 may result in an increase in intracellular calcium ion levels, subsequently promoting actin polymerization. This may, in turn, bolster the integrity of AIS actin rings during homeostatic structural plasticity.

## Conclusion

Our study aimed to examine the changes in AIS along the axon and the periodicity of actin rings at the AIS during homeostatic structural neuronal plasticity. We achieved this by applying different drugs to either depolarize or hyperpolarize the plasma membrane. Using confocal and STED super-resolution microscopy, we assessed the localization of the AIS and the periodicity of actin rings at the AIS in immature and mature neurons. Our findings revealed that incubation of neurons with the Kv7 channel blocker XE991 undermined the integrity of actin rings in immature neurons at the AIS at DIV10 but had no such effect in mature neurons at DIV16. In contrast, both the Nav channel blocker TTX and the Kv7 channel opener RTG significantly decreased the integrity of actin rings at the AIS at both DIVs. We hypothesize that prolonged exposure to TTX possibly diminished the crowding of the plasma membrane of the AIS in mature neurons. Additionally, long-term incubation with TTX and led to depolymerization of actin rings at the AIS. Furthermore, TTX and RTG might also have affected the integrity of actin rings through decreasing intracellular calcium ion levels by partially blocking calcium influx. Additional work is needed to elucidate the specific molecular mechanisms that underlie these various forms of homeostatic structural neuronal plasticity.

## Author Contributions

S. Gu, A.V. Tzingounis, and G. Lykotrafitis designed the research; S. Gu conducted the research and collected the data; S. Gu, A.V. Tzingounis, and G. Lykotrafitis analyzed and interpreted the data; A.V. Tzingounis and G. Lykotrafitis contributed new reagents and analytic tools; S. Gu, A.V. Tzingounis, and G. Lykotrafitis wrote the paper.

## Declaration of interests

The authors declare no conflicts of interest.

## Acknowledgments

This work was supported by the National Science Foundation Division of Physics, Physics of Living Systems 2210535 to G. Lykotrafitis. The Abberior STED microscope was acquired through funding from the NIH grant S10OD023618, which was awarded to Christopher O’Connell.

## Data Availability

The data of this article are available from the corresponding author upon reasonable request.

## Supporting material

**Figure S1.**
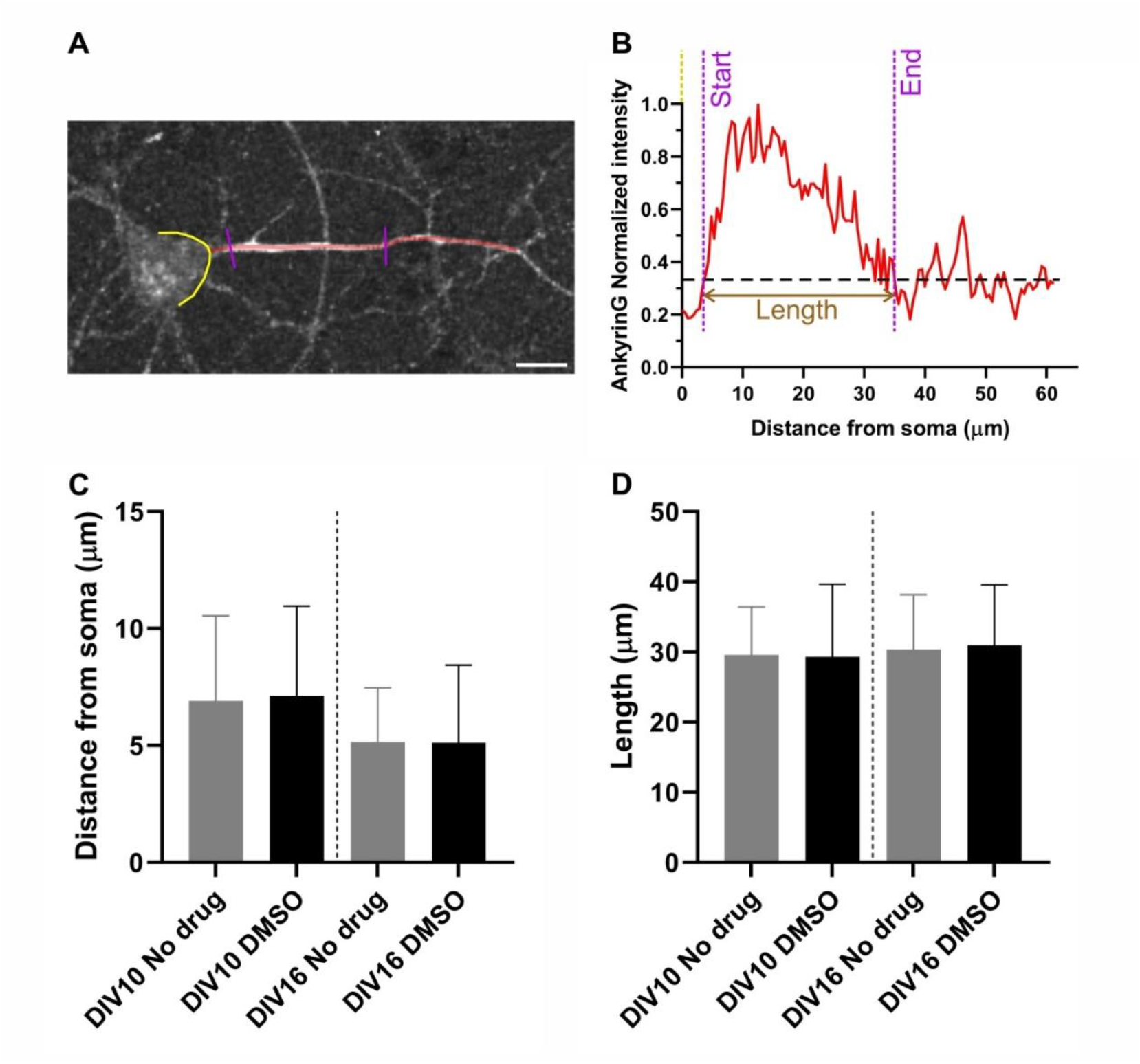
DMSO treatment does not alter the distribution of ankG along the axon. *(A)* Representative confocal image of ankG immunostaining along the axon. Scale bar, 10 µm. *(B)* Normalized fluorescence intensity of ankG corresponding to the red line region in *(A)*. The black dashed line indicates the 0.33 threshold, which is used to define the start and end locations of ankG. The yellow dashed line indicates the edge of the soma. The magenta dashed line denotes the determined start and end positions of ankG and the brown line represents the length of ankG. *(C)* Plots showing the comparison of the start position of ankG along the axon at DIV10 and DIV16 between no drugs treated and DMSO treated group. Sample size: DIV10 No drug: n = 23; DIV10 DMSO: n = 23; DIV16 No drug: n = 22; DIV16 DMSO: n = 25. *(D)* Plots showing the comparison of the length of ankG along the axon at DIV10 and DIV16 between no drugs treated and DMSO treated group. Sample size: DIV10 No drug: n = 23; DIV10 DMSO: n = 23; DIV16 No drug: n = 22; DIV16 DMSO: n = 25. Non-parametric Mann-Whitney tests were performed. All plots represent the mean ± SD.

**Figure S2.**
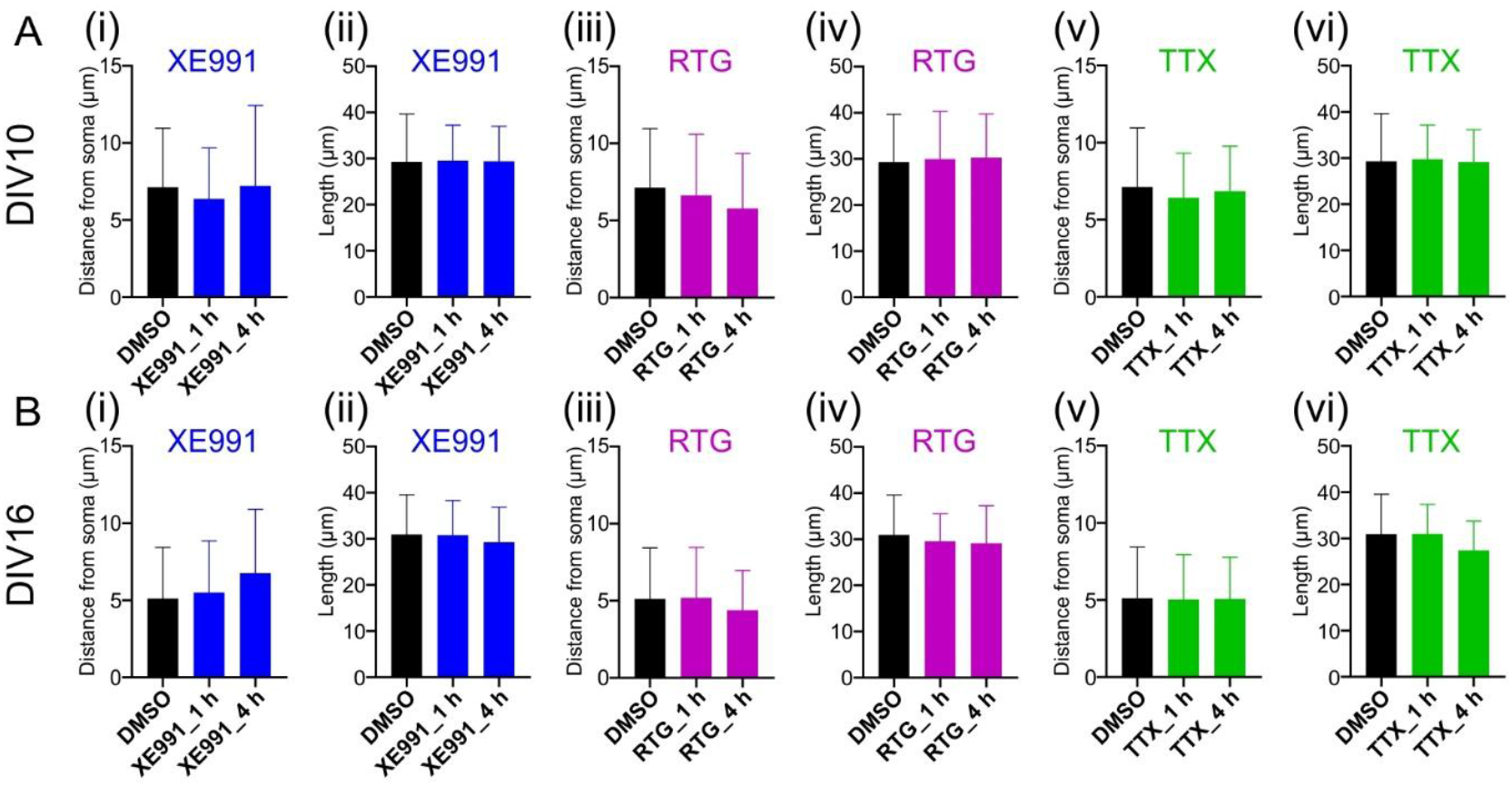
Short-term treatment with XE991, RTG, or TTX does not significantly alter the distribution of ankG along the axon. *(A)* Plots showing the start position and length of ankG along the axon at DIV10 after treatment with XE991 *(i-ii)*, RTG *(iii-iv)*, or TTX *(v-vi)* for 1 and 4 h. Sample size: *(i)* and *(ii)*: DMSO: n = 23; XE991_1 h: n = 18; XE991_4 h: n = 18. *(iii)* and *(iv)*: DMSO: n = 23; RTG_1 h: n = 18; RTG_4 h: n = 22. *(v)* and *(vi)*: DMSO: n = 23; TTX_1 h: n = 18; TTX_4 h: n = 23. *(B)* Plots showing the start position and length of ankG along the axon at DIV16 following the same treatments as applied at DIV10 in *(A)*. Sample size: *(i)* and *(ii)*: DMSO: n = 25; XE991_1 h: n = 23; XE991_4 h: n = 23. *(iii)* and *(iv)*: DMSO: n = 25; RTG_1 h: n = 20; RTG_4 h: n = 22. *(v)* and *(vi)*: DMSO: n = 25; TTX_1 h: n = 21; TTX_4 h: n = 21. Non-parametric one-way ANOVA test was performed. All plots represent the mean ± SD.

**Figure S3.**
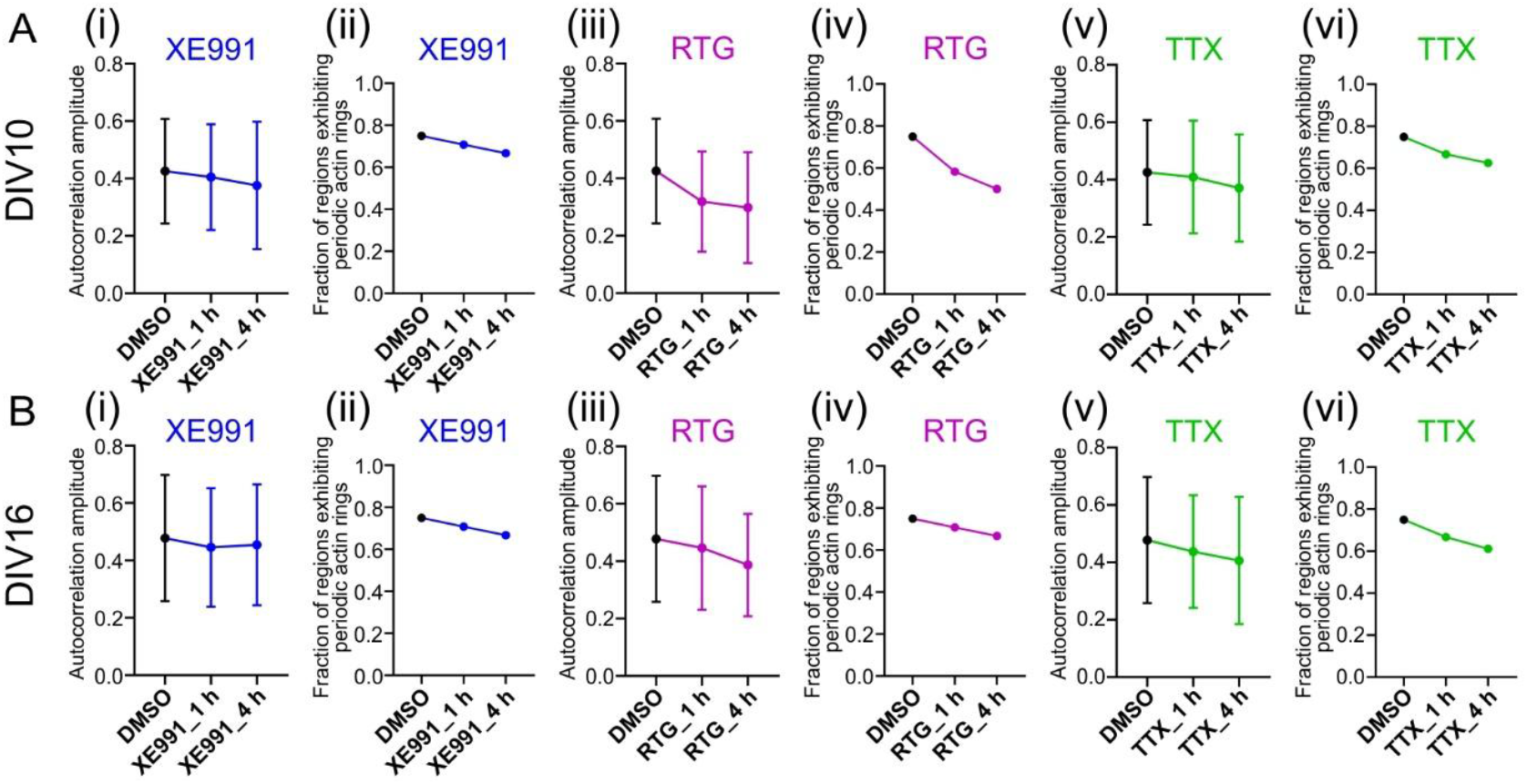
Short-term treatment with XE991, RTG, or TTX does not significantly alter the periodicity of actin rings at the AIS. (*A*) Plots showing autocorrelation amplitude and fraction of regions exhibiting periodic actin rings following treatment at DIV10 with XE991 *(i-ii)*, RTG *(iii-iv)* or TTX *(v-vi)*. Sample size: n = 24 for each treatment group. *(B)* Plots showing autocorrelation amplitude and fraction of regions exhibiting periodic actin rings at DIV16 following the same treatments at DIV10 in *(A)*. Sample size: n = 24 for each treatment group. Non-parametric one-way ANOVA tests were performed. All plots represent the mean ± SD.

## Notes

### Competing Interest Statement

The authors have declared no competing interest.

